# Neutralizing antibody responses to an HIV envelope glycan hole are not easily broadened

**DOI:** 10.1101/827477

**Authors:** Yuhe R. Yang, Laura E. McCoy, Marit J. van Gils, Raiees Andrabi, Hannah L. Turner, Meng Yuan, Christopher A. Cottrell, Gabriel Ozorowski, James Voss, Matthias Pauthner, Thomas M. Polveroni, Terrence Messmer, Ian A. Wilson, Rogier W. Sanders, Dennis R. Burton, Andrew B. Ward

**Affiliations:** Department of Integrative Structural and Computational Biology and the Skaggs Institute for Chemical Biology, The Scripps Research Institute, La Jolla, CA 92037, USA; Scripps Consortium for HIV/AIDS Vaccine Development (CHAVD), The Scripps Research Institute, La Jolla, CA 92037, USA; IAVI Neutralizing Antibody Center and the Collaboration for AIDS Vaccine Discovery (CAVD), The Scripps Research Institute, La Jolla, CA 92037, USA; Division of Infection and Immunity, University College London, London, UK; Department of Medical Microbiology, Amsterdam UMC, University of Amsterdam, 1105 AZ Amsterdam, the Netherlands; Department of Immunology and Microbiology, The Scripps Research Institute, La Jolla, CA 92037, USA; Scripps Consortium for HIV/AIDS Vaccine Development (CHAVD), The Scripps Research Institute, La Jolla, CA 92037, USA; IAVI Neutralizing Antibody Center and the Collaboration for AIDS Vaccine Discovery (CAVD), The Scripps Research Institute, La Jolla, CA 92037, USA; Skaggs Institute for Chemical Biology, The Scripps Research Institute, La Jolla, CA 92037; Department of Microbiology and Immunology, Weill Medical College of Cornell University, New York, NY 10065, USA; Ragon Institute of Massachusetts General Hospital, Massachusetts Institute of Technology, and Harvard University, Cambridge, MA 02139, USA

**Keywords:** HIV-1, SOSIP, glycan hole, epitope, rabbit immunization, monoclonal antibodies, autologous neutralization

## Abstract

Extensive studies with subtype A BG505-derived HIV envelope glycoprotein (Env) SOSIP immunogens have revealed that the dominant autologous neutralizing site in rabbits is located in an exposed region of the heavily glycosylated trimer that lacks potential N-linked glycosylation sites at positions 230, 241, and 289. The Env derived from B41, a subtype B virus, shares a glycan hole centered on positions 230 and 289. BG505 and B41 SOSIP immunogens were combined to test whether immunization in rabbits could induce broader Tier 2 neutralizing responses to the common glycan hole shared between BG505 and B41. Here we isolated autologous neutralizing antibodies (nAbs) that were induced by immunization with B41 SOSIP alone, as well as B41 and BG505 co-immunization, and describe their structure in complex with the B41 SOSIP trimer. Our data suggest that distinct autologous nAb lineages are induced by BG505 and B41 immunogens, even when both immunogens were administered together. In contrast to previously described BG505 glycan hole antibodies, the B41-specific nAbs accommodate the highly conserved N241 glycan (>97% conserved), which is present in B41. Single particle cryo-electron microscopy (cryoEM) studies confirmed that B41 and BG505-specific nAbs bind to overlapping glycan hole epitopes. In an attempt to broaden the reactivity of a B41-specific nAb, mutations in the BG505 glycan hole epitope guided by our high-resolution data only recovered partial binding. Overall, designing prime-boost immunogens to increase the breath of nAb responses directed at glycan holes epitopes remains challenging even when the typically immunodominant glycan holes despite overlap with different Envs.

**IMPORTANCE:** A glycan hole is one of the most dominant autologous neutralizing epitopes targeted on BG505 and B41 SOSIP trimer immunized rabbits. Our high-resolution cryoEM studies of B41 in complex with a B41-specific antibody complex elucidate the molecular basis of this strain-specific glycan hole response. We conclude that eliciting cross-reactive responses to this region would likely require hybrid immunogens that bridge between BG505 and B41.

## INTRODUCTION

With ∼1.7 million new infections in 2018, human immunodeficiency virus (HIV) continues to be a major global public health issue (data from http://aidsinfo.unaids.org/). Although antiretroviral therapies (ARTs) have dramatically reduced mortality, preventative vaccines would be invaluable to control the spread of the pathogen. The human antibody response to HIV envelope glycoprotein (Env) following infection predominantly binds non-fusogenic conformations of Env, often referred to as “viral debris”, as opposed to the intact fusogenic form displayed on the surface of the virus (1, 2). The corresponding antibodies are termed non-neutralizing and often recognize epitopes displayed both by conformationally open or partially disassembled Env and by glycoprotein 41 (gp41) subunit stumps, which remain after the glycoprotein 120 (gp120) subunit dissociates from the Env trimer. Infection can also elicit functional antibodies that bind the intact Env trimer and neutralize the virus strain prevalent in the infected host (3, 4). However, the virus can rapidly escape these strain-specific neutralizing antibodies (nAbs) by mutating the sequence within and surrounding the epitope and by adding glycosylation sites (5). In contrast, a small proportion of HIV-infected individuals develop broadly neutralizing antibodies (bnAbs), which recognize epitopes comprised of relatively conserved amino acids as well as N-linked glycans (6-8). These bnAbs are capable of neutralizing a high percentage of HIV strains and their development has been associated with longer exposure to multiple evolving strains of the HIV virus (8). One approach to develop an effective vaccine capable of bnAb elicitation therefore involves cocktails of different trimer immunogens as well as sequential immunization with Envs derived from different strains (9, 10).

Antibodies elicited against stabilized HIV Env immunogen trimers (11, 12) can exhibit robust neutralization against immunogen-matched neutralization resistant (Tier 2) viruses in animal models. Our previous work with BG505-derived immunogens in rabbits has revealed that an autologous neutralizing epitope region on BG505 is exposed and immunodominant due to the absence of glycan sites at positions 241, which is present in >97% of HIV strains, at position 289, which is present in >70% HIV strains, and at position 230, which is less conserved and only present in <35% of HIV strains (13-16). Certain other HIV strains lack some of the same glycan sites in the Env as BG505, resulting in partially overlapping holes in their glycan shield. These findings raised the question whether Env immunogens with overlapping glycan holes could be combined to induce broader Tier 2 neutralizing responses. A stabilized SOSIP.664 trimer derived from a subtype B Env gene, named B41, has been described previously (17) and has a partially overlapping glycan hole with that of BG505. Like BG505, B41 SOSIP lacks glycans at positions 289 and 230, but does contains the more highly conserved glycan at 241. Previous immunization studies with B41 SOSIP revealed that the majority of nAb response in rabbits was indeed directed to the 289 glycan hole (17). However, the exact epitope and molecular details of the interactions with nAbs remain unknown.

In this study, we isolated B41-specific monoclonal Abs (mAbs) and confirmed that the dominant B41 autologous neutralizing response targets the 230/289 glycan hole on the B41 immunogen. Importantly, these nAbs can accommodate the highly conserved N241 glycan. Even when both BG505 and B41 immunogens were administered together, the isolated B41-specific nAb lineages were unable to cross-neutralize BG505 indicating no, or very limited, cross-boosting. We also show the molecular details of a B41-specific nAb bound to the 230/289 glycan hole epitope using high-resolution cryo-electron microscopy (cryoEM). Based on the amino-acid contact residues between B41 and a B41-specific nAb, we then mutated BG505 to gain some binding of the B41 strain-specific nAb. In summary, we established that B41 and BG505-specific nAbs recognize different amino acids in their corresponding epitopes to block viral infection and identified key residues that contribute to the antibody specificity. While our B-cell isolation and antibody production was by no means exhaustive, given the prevalence of nAbs elicited against the shared glycan hole epitope in BG505 and B41, it is notable that no cross-nAbs were isolated. Interestingly, a large number of the non-neutralizing antibodies isolated were cross-reactive, but likely bind to the irrelevant trimer base epitope. Therefore, we conclude that designing prime-boost or cocktail immunization regimens that increase the breath of glycan hole directed nAb responses remains a challenge even with immunogens that share glycan holes.

## RESULTS

### B41 Env trimers induce autologous nAbs that do not cross-neutralize BG505

Four rabbits from a previously described immunization experiment (14) were used to isolate mAbs for the current study. In the prior study, animals were separated into two groups: group 1 (5713 & 5716) received 30 µg B41 SOSIP trimer alone per immunization (large arrows), while group 2 (5746 & 5749) received a bivalent cocktail containing both BG505 SOSIP and B41 SOSIP in a 1:1 ratio (10 µg or 30µg per immunization, small and large arrows, respectively) (FIG 1A). Group 1 animals 5713 and 5716, which only received the B41 immunogen, both had ID_50_ neutralization titers against the wildtype B41 pseudovirus of around 1 in 700 (FIG 1B), as expected given the single immunogen used. Of the two animals that received both immunogens only 5746 has cross-neutralizing sera. Rabbit 5746, which received a low dose of the BG505/B41cocktail, had a higher neutralizing titer (∼3300) against the wildtype B41 pseudovirus than BG505 pseudovirus (∼1800) (FIG 1B). Rabbit 5749, which received a high dose of the BG505 SOSIP and B41 SOSIP cocktail, is BG505-specific and undetectable neutralizing titer against the wildtype B41 pseudovirus (FIG 1B).

**FIG 1.**
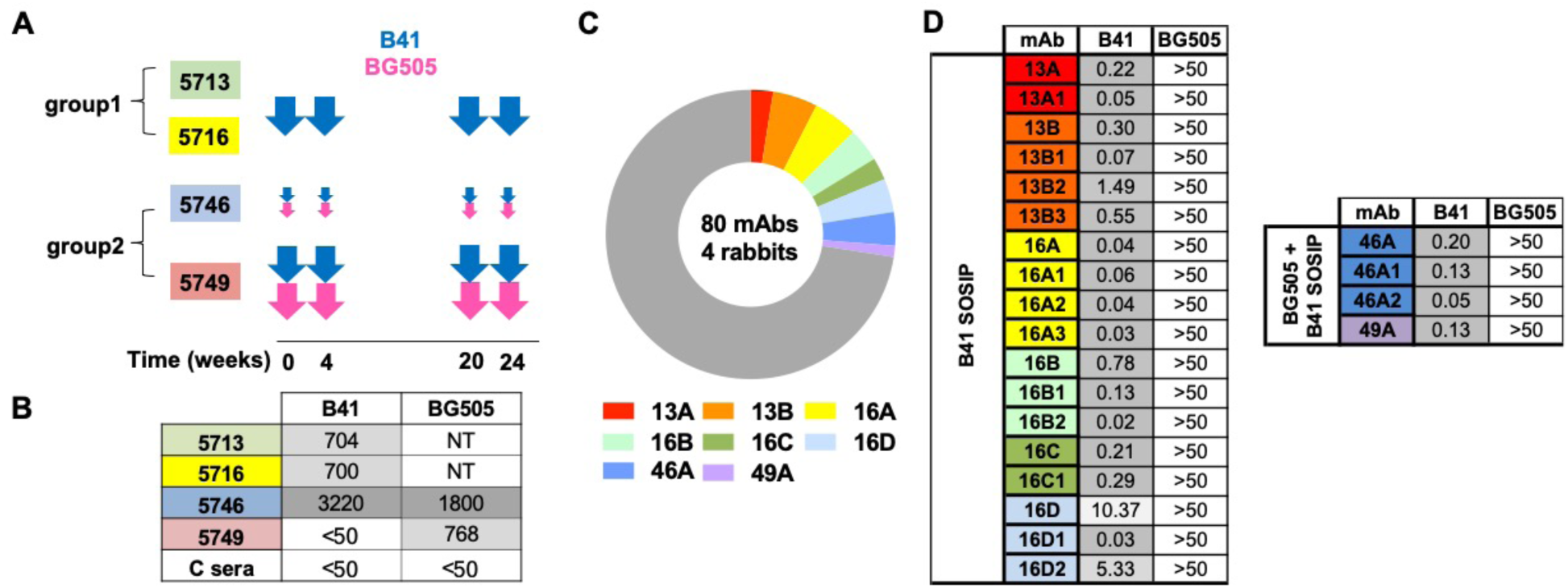
Immunization schedule and autologous neutralizing activities of B41-specific rabbit mAbs. (A) Schematic of immunization schedule of four individual rabbits from two groups. Each immunization is indicated by an arrow and every animal was immunized at the listed timepoints as described in Klasse *et al.* (14). (B) Neutralization titers (ID_50_) of immunized rabbit sera against B41 and BG505 pseudoviruses. C sera is from an unimmunized rabbit 3421 described in the previous study (13) as a control. (C) Pie chart showing that 22 of the 80 mAbs (28%) derived from four rabbits can neutralize the immunogen-matched B41 pseudovirus (rabbit mAb families color-coded as in the legend below). (D) Neutralization analysis of isolated nAbs against B41 and BG505 pseudoviruses. Inhibitory concentration (IC_50_) values in µg/ml are listed.

From these four animals, 80 mAbs that bound B41 SOSIP were isolated by single B cell sorting using B41 SOSIP as the bait and successfully PCR amplified (FIG 1C). MAbs were named with a similar nomenclature as described previously (13). Each mAb was named with the rabbit identifier (13, 16, 46 & 49) and then a unique alphabetical lineage identifier (e.g. A, B etc). Lineage members were then assigned with an additional number: 13A1, 13B1, 13B2 etc. 58 mAbs (72%) bound both BG505 and B41 immunogens but were unable to neutralize either BG505 or B41 pseudoviruses. Although these mAbs were not studied further, we suspect that the majority of them target the immunodominant epitope present on the soluble SOSIP trimer but not the viral surface membrane embedded trimer (18). In contrast, 22 mAbs (28%) bound only the B41 immunogen. These mAbs derived from eight genetically distinct families and all family members were able to neutralize the immunogen-matched B41 pseudovirus (FIG 1C). The B41-specific nAbs exhibited strong neutralization against B41 with IC_50_ values as low as 0.02 µg/ml (FIG 1D). Interestingly, none of these nAbs showed cross neutralization of BG505, even the mAbs from the 46A and 49A families isolated from the rabbits that received the BG505 and B41 SOSIP trimer cocktail. These data suggest that the BG505 and B41 immunogens appear to induce independent autologous nAb responses.

### B41-specific rabbit nAbs target the 230/289 glycan hole

Our previous work with BG505 immunogens in rabbits revealed that the dominant autologous nAbs target the glycan hole created by the absence of glycans at positions 230, 241, and 289 (13). Moreover, autologous neutralization specific for glycan holes was also seen following immunization with trimers from different clades (19). To determine whether the B41 neutralizing mAbs also targeted a glycan hole on the B41 immunogen, we tested the neutralization activity of eight isolated mAbs representing the different autologous nAb families using a panel of B41 mutant pseudoviruses with the N230 and N289 glycans knocked-in (FIG 2A&B). Introduction of the N289 glycan abolished or greatly reduced the neutralization activity for all eight mAbs (FIG 2A), which was reflected in a significant reduction of the maximum neutralization capacity (FIG 2B). The detrimental effect of introducing the N289 glycan was largely mitigated when the pseudovirus was grown in the presence of kifunensine to enrich for oligomannose glycans (FIG 2A&B), where the maximum neutralization values were increased above the 50% neutralization threshold allowing the calculation of IC_50_ values. In contrast, neutralization activity for all mAbs was only mildly affected when the wildtype B41 pseudovirus was grown in the presence of kifunensine. Furthermore, while introduction of the N230 glycan (with or without kifunensine) eliminated neutralization activity for mAbs 13B and 49A, it had no effect on the other six mAbs. The role of the N241 glycan was tested with B41 N241-knock out (KO) pseudovirus, and showed that neutralization was in some cases diminished, but not abolished entirely for any of the mAbs.

**FIG 2.**
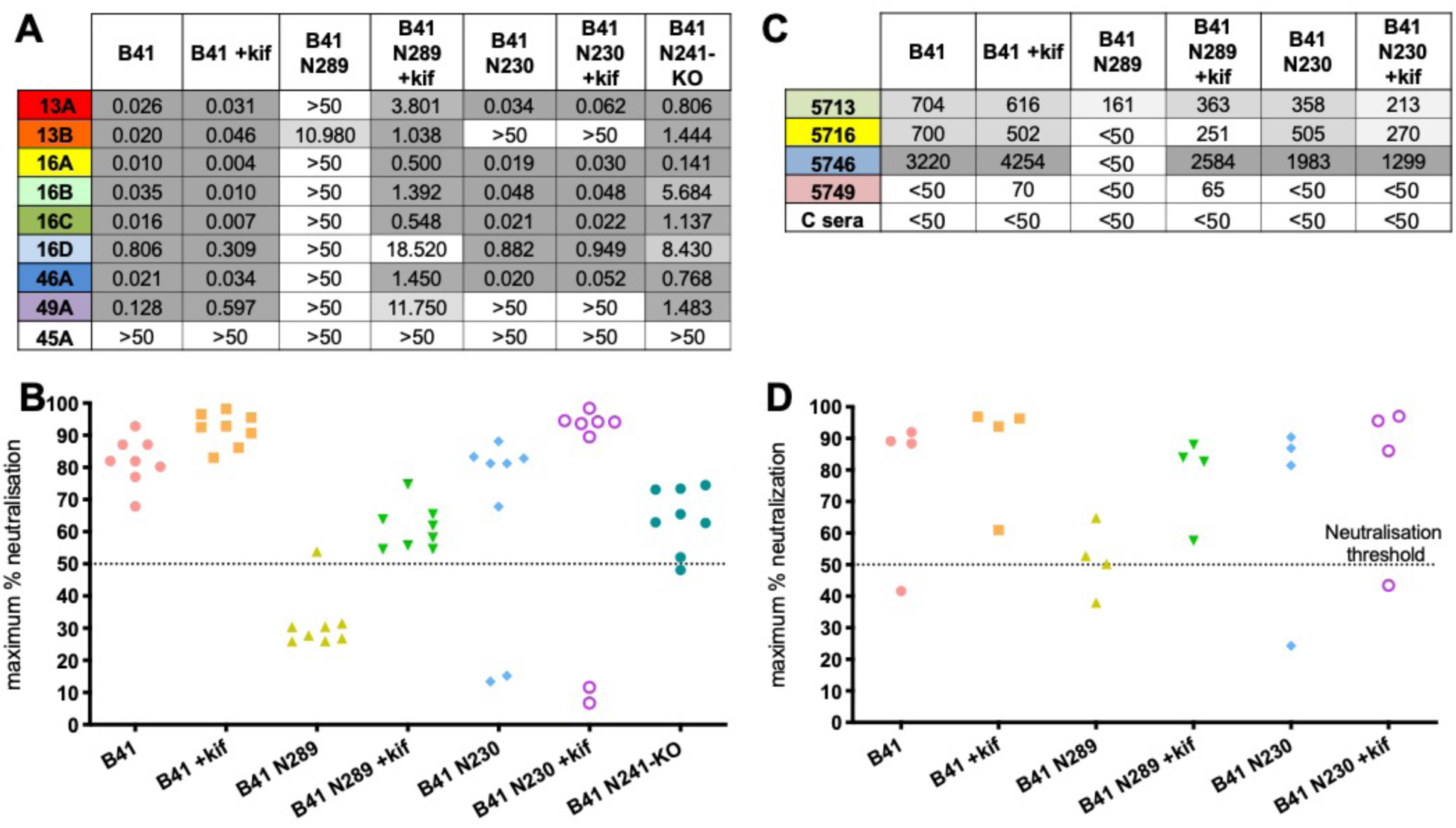
B41-specific rabbit nAbs target the 230/289 glycan hole. (A) Neutralization analysis of isolated nAbs against a panel of B41 mutants covering glycans at site 230, 289 and 241. +kif indicates that the pseudovirus was grown in the presence of kifunensine. Inhibitory concentration (IC_50_) values are listed in µg/ml. Non-nAb 45A was used as a control. The highest concentration tested for neutralization was 50 µg/ml. (B) Maximum neutralization percentage of isolated nAbs against autologous B41 virus and mutants. (C) Neutralization titers (ID_50_) of immunized rabbit sera determined using B41 mutants. C sera is from an unimmunized rabbit 3421 as a control. NT stands for non-tested. (D) Maximum neutralization percentage of sera against autologous B41 virus and mutants. The Y axis shows the max neutralization percentage for sera from individual rabbits.

We then tested sera neutralization activity with the same panel of B41 mutant pseudoviruses to evaluate if the activity of individual mAbs is representative of the activity in the sera (FIG 2C&D). Consistent with the observations made using the mAbs, the introduction of a glycan site at position 289 greatly decreased neutralization activity in all 4 rabbit sera (FIG 2C&D), suggesting that the isolated mAbs represent a substantial proportion of the nAbs within the sera. Moreover, the same restoration of neutralization activity was observed when the sera were tested against the N289-KI virus expressed in the presence of kifunensine. In addition, the introduction of a glycan site at position 230 had relatively little effect on serum neutralization activity.

### B41-specific rabbit nAbs resemble BG505-specific glycan hole nAbs

Enzyme-linked immunosorbent assay (ELISA) binding assays showed that all eight mAbs bound to B41 SOSIP, while 6/8 also bound to B41 gp120 (Fig 3A). MAbs 13B and 49A were not able to bind to B41 gp120 (Fig 3A). Given the gp120-specific nature of their epitopes revealed by our structural studies (below) the most likely reason for not binding is a difference in the glycosylation pattern of gp120 versus the Env trimer. Non-nAbs 45A and 48A, which were isolated in parallel from a previous study (14), show strong binding to B41 SOSIP trimer but not to gp120. Competition ELISAs using previously described bnAbs that target distinct epitopes were conducted to the B41 mAbs (Fig 3B &S5). PG9, PGT121, 8ANC195, and PGV04, that target the trimer apex, N332-glycan supersite, gp120-gp41 interface, and CD4-binding site epitopes, respectively, were used in the analysis (Fig 3C). The results showed that 49A and 13B compete with the human gp120-gp41 interface specific bnAb 8ANC195, indicative of overlapping epitopes. The B41 nAbs also exhibited a high level of competition between themselves suggesting that they targeted a common epitope (FIG S5).

**FIG 3.**
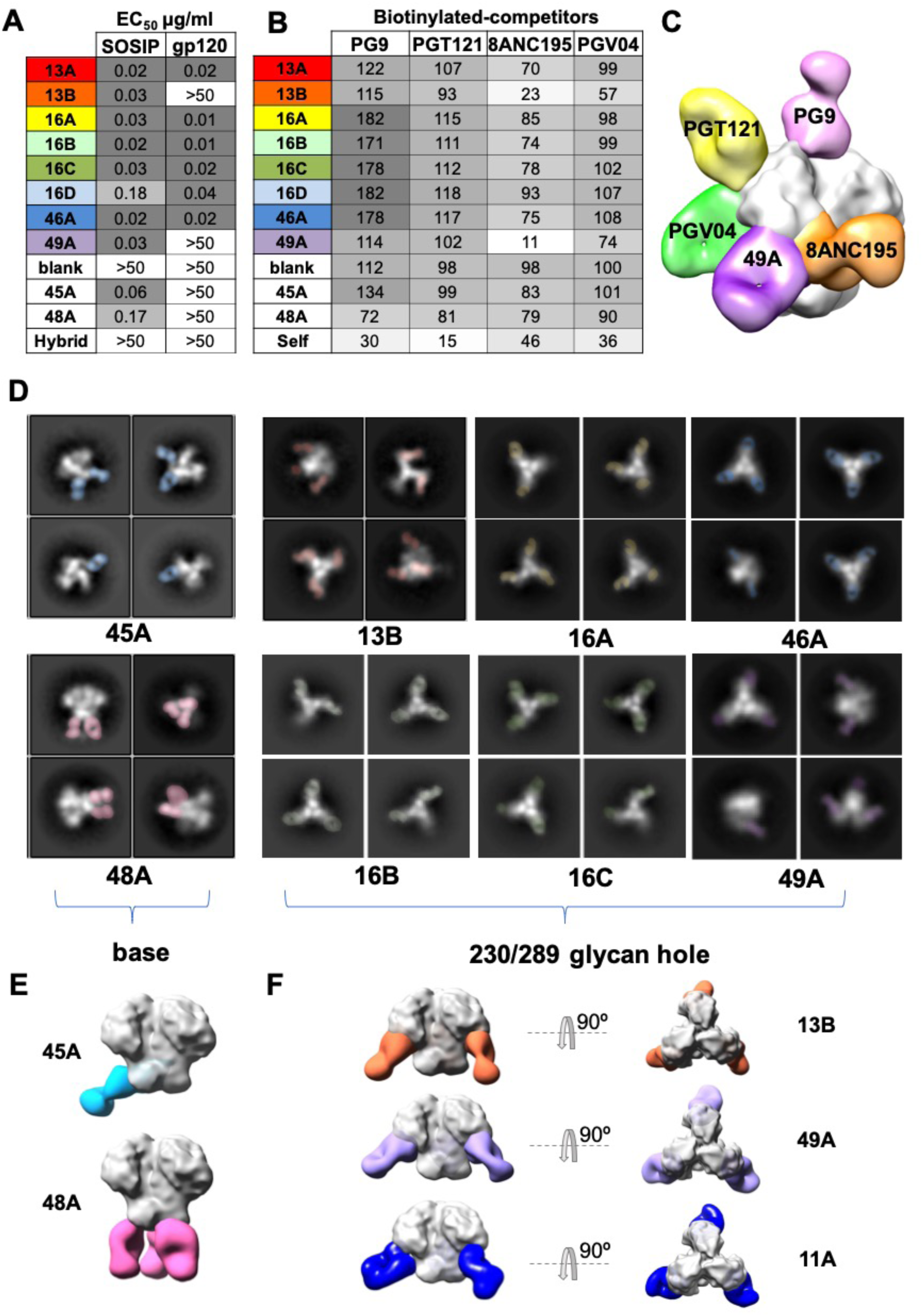
Epitope mapping by competition assay and negative-stain electron microscopy (NS-EM). (A) Enzyme-linked immunosorbent assay (ELISA) of isolated mAbs against the B41 gp120 monomer and B41 SOSIP trimer. The half-maximal effective concentrations (EC_50_) in µg/ml are listed. Non-nAbs (45A and 48A) were isolated in parallel with the antibodies described here. The negative control mAb, named hybrid, was made from the heavy chain of R56 and light chain of R20 (PDB: 4JO3) used in a previous study (13). (B) Competition ELISAs of isolated mAbs against previously identified bnAbs. The percent binding of biotinylated bnAbs was tested in the presence of the indicated non-biotinylated rabbit mAb competitors. The data represent the percentage reciprocal binding where 100% was the absorbance measured for each bnAb in the absence of any competitor. (C) 3D reconstruction comparison of nAb 49A epitope to previously identified bnAbs. (D) Representative 2D classes (bottom) of different mAbs. (E) Representative 3D reconstructions of base binding antibodies, 45A (blue) and 48A (pink), and (F) glycan hole targeting antibodies including 13B (orange), 49A (purple) and 11A (dark blue) bound to B41 SOSIP.

We next carried out single particle negative-stain electron microscopy (NS-EM) to more precisely determine the location of the epitopes targeted by the isolated mAbs. Epitope mapping of 8 mAbs resulted in only two classes of epitopes (FIG 3D-F). Class 1 contained the non-neutralizing base binders, 45A and 48A, that targeted the base of the B41 trimer at different angles (FIG 3E &S2). Representative 2D classes showed that 48A bind with a stoichiometry of three antibodies per trimer while 45A only bind with one to two antibodies per trimer. Class 2 included 6 nAbs targeting an overlapping epitope around the 230/289 glycan hole region, confirming the mutant neutralization and competition binding results. Representative 2D classes showed that all nAbs bind with a stoichiometry of three antibodies per trimer. Representative 3D EM reconstructions from B41-13B and B41-49A complexes were shown to further illustrate the epitope (FIG 3F). The epitope overlaps considerably with the BG505 specific glycan hole antibody 11A, and the antibodies approach the trimer with similar upward angles (FIG 3F).

NS-EM, ELISA and neutralization data confirm that the B41-specific antibodies target a similar glycan hole region as the previously described BG505-specific mAbs 10A, 11A and 11B (13). However, our neutralization results demonstrate that B41 isolated mAbs lack the ability to neutralize BG505 (FIG 1D), including the 46A and 49A family nAbs that are elicited in rabbits immunized with the B41 and BG505 cocktail. To further understand why there was no cross-reactivity of neutralizing Abs targeting an overlapping glycan hole epitope, cryoEM structural studies were conducted on B41 SOSIP in complex with 13B. While 13B was isolated from a B41 SOSIP only immunized animal it was representative of all the B41-specific nAbs as illustrated by the structural similarity revealed by NS-EM.

### A high-resolution cryoEM structure of 13B in complex with the B41 SOSIP trimer reveals atomic details of recognition

To characterize the binding mode of the B41-specific antibodies, we have obtained a ∼3.9 Å resolution cryo-EM map reconstruction of nAb 13B in complex with the B41 SOSIP trimers and built an atomic model (FIG. 4A). A starting model was created by combining a homology model of the Fv region of 13B generated using the Rosetta antibody protocol (20) and a B41 SOSIP crystal structure (PDB 6MCO). 13B binds to the glycan hole epitope with the heavy chain making the majority of contacts. This is different from the BG505 glycan hole mAbs which appear to interact primarily via the light chain (18). The glycan hole epitope is surrounded by 8 glycans including N88, N234, N241, N276, N295, N339, N355 and N448 (FIG. 4B), which likely constrain the angle of approach for the elicited antibodies. The first two sugars of the N241 glycan (which is present in B41 but not BG505) were resolved in the refined map (FIG. 4C). The glycan density is in close proximity to the 13B density, but remains distinct, indicating no direct contact. The identical conformation for the N241 glycan was observed in our B41-13B structure as well as a previously solved crystal structure of B41 (PDB 6MCO), indicating that the binding of 13B did not result in a conformational change of the glycan. These data are consistent with our conclusion that 13B does not specifically interact with the N241 glycan.

**FIG. 4.**
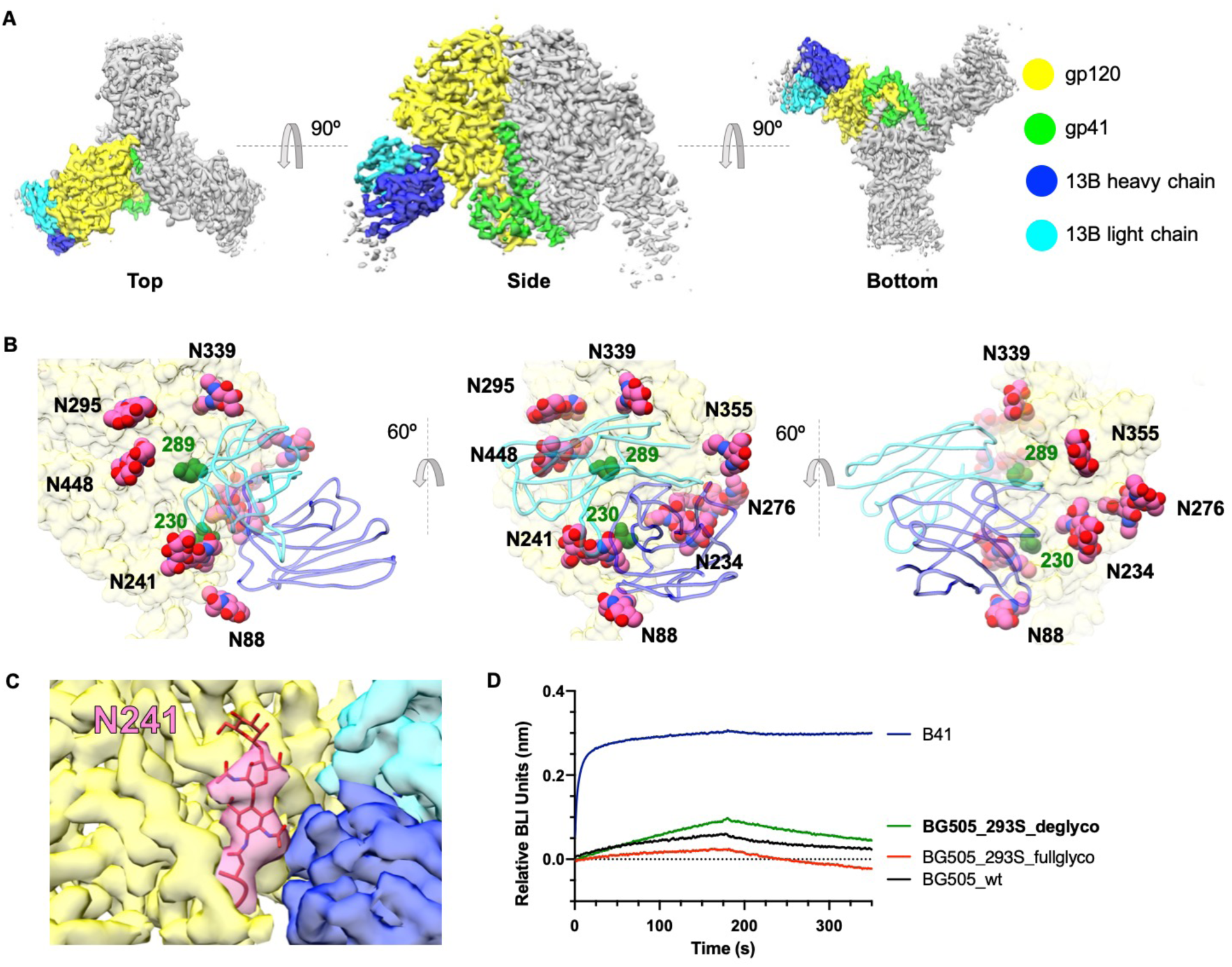
CryoEM map and model of B41-13B at 3.9 Å. (A) Top, side and bottom view of a cryoEM 3D reconstruction of B41-13B complex at ∼3.9 Å resolution colored by subunits. (B) Zoomed in image showing how antibody13B targets the 230/289 glycan hole epitope (230 and 289 residues highlighted in dark green); modeled glycans (spheres) in pink, ribbon representation of 13B Fab in blue (dark: heavy chain, light: light chain) and B41 gp120 surface in yellow. (C) The density map shows that 13B (light and dark blue) interacts exclusively with gp120 peptide (yellow) and although the N241 glycan (highlighted in pink) is close by no direct contacts are observed. When the previously solved crystal structure (PDB 6MCO) of B41 SOSIP was docked in the N241 glycan (red sticks) fit exactly into the cryoEM density for the glycan. (D) Biolayer interferometry (BLI) binding sensorgrams showing association and dissociation of His-tagged B41 SOSIP (blue) and BG505 SOSIP variants with 13B Fab. Deglycosylated BG505 SOSIP is colored in green, HEK293S-expressed BG505_fullglyco without EndoH treatment in red and HEK293F-expressed BG505_wt SOSIP in black.

To further evaluate the role of glycans in antibody binding, a deglycosylated version of BG505 SOSIP was prepared by expression in HEK293S cells followed by EndoH treatment (FIG S3A). Biolayer interferometry (BLI) analysis against 13B was compared in parallel with positive control B41, negative control wild type BG505 as well as HEK293S-expressed BG505 before EndoH treatment (FIG. 4D). The removal of glycans of BG505 only resulted in a subtle impact on binding to 13B, and differences in binding affinity could not be reliably calculated with the observed binding curves. When we further tested binding of 13B with deglycosylated BG505 by NS-EM, no complex formation was observed (FIG S3B). These data support the conclusion that the lack of cross-reactivity to BG505 is therefore not due glycan differences. Thus, we concluded that protein sequence differences between BG505 and B41 within the epitope region are responsible for the lack of cross reactivity.

The high resolution structure of B41 allowed direct comparison of the epitope regions of B41-specific and BG505-elicited rabbit mAbs (13). In order to identify residues in B41 that contribute to binding, we highlighted all of the potential contact residues of B41 and 13B (FIG 5A). The contact residues are defined as two residues containing any atom within 4 Å of each other, as determined using Chimera (21). Next, we superimposed the high-resolution BG505 model (PDB:5CEZ) on the gp120 subunit of B41 SOSIP to identifiy potential clashing residues, which we defined as atoms closer than 1.0 Å (FIG 5B). The potential clashes involved three residues in the heavy chain: Y98 (heavy chain) with K232 (Env), P100_B_ and 100S with Q348, and P100_B_ with K351, as well as one in the light chain: R95 with N355. Four residues in the epitope regions differ between B41 and BG505, as shown in the sequence alignment in the table (FIG. 5C). Interestingly, despite being identical between the two Env residues 268 and 269 clash in our BG505 docked model, indicating that neighboring residues that differ between the strains caused structural perturbations in these conserved residues.

**FIG 5.**
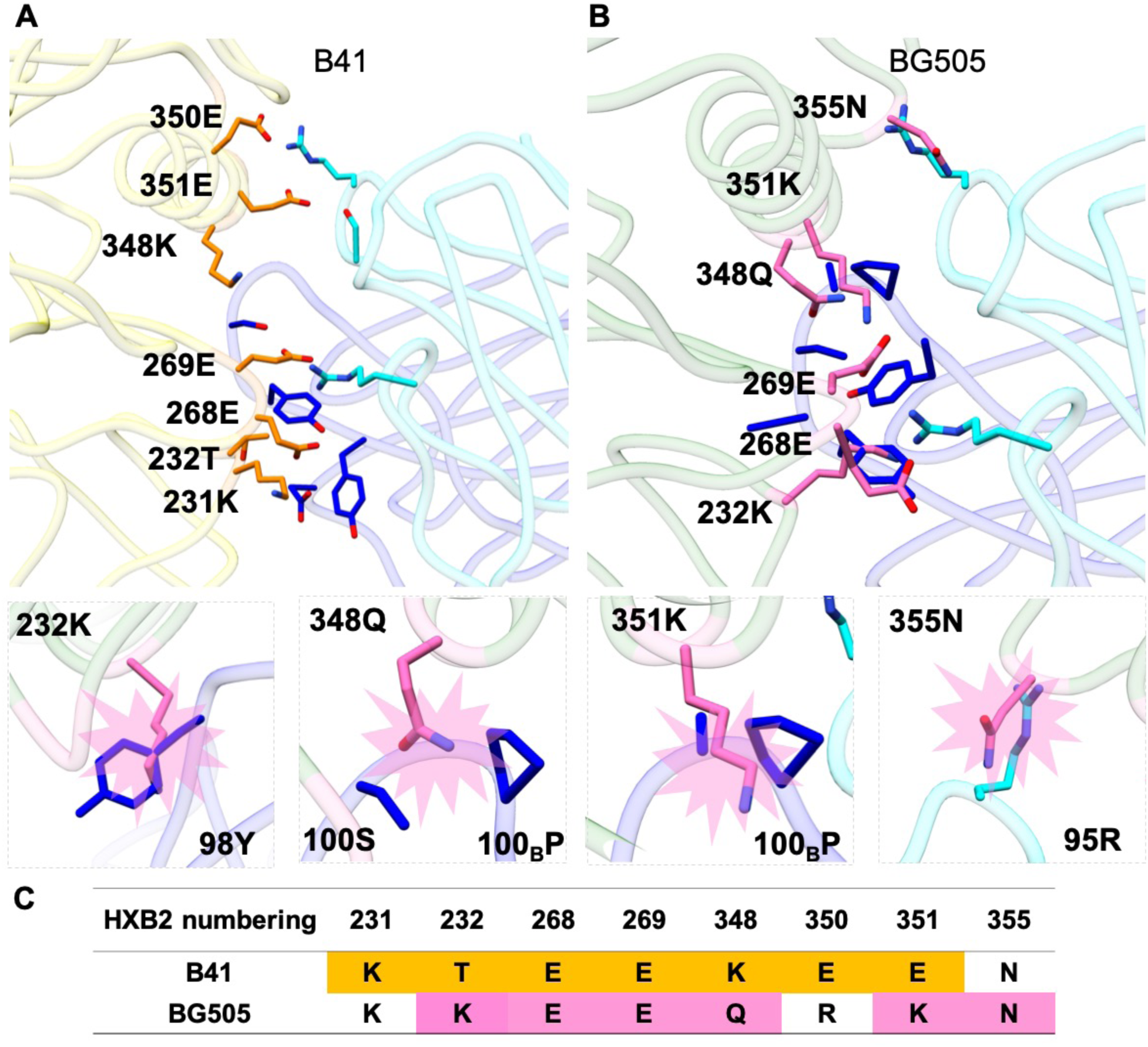
Structural comparison of BG505 SOSIP and B41 SOSIP with B41-elicited rabbit antibody 13B bound to its epitope. (A) Contact residues at the B41-13B interface. Contact residues are defined as two residues containing any atom within 4 Å of each other; B41 residues are colored orange and 13B residues in light (light chain) and dark (heavy chain) blue. (B) Superimposition of BG505 (PDB:5CEZ) and B41-13B complex aligned on gp120. For clarity, the B41 trimer is not shown. Potential residues that clash between BG505 and 13B are highlighted in pink sticks. Below are zoomed-in structures of 4 potential clashes of BG505 with 13B involving three with the heavy chain (232K-98Y, 348Q-100_B_P//100S, 351K-100_B_P) and one with the light chain (355N-95R). The heavy chain is in dark blue, BG505 potential clashing residues in pink, light chain residues in light blue, and gp120 of BG505 in light green.(C) Sequence alignment of potential contact residues of B41 (highlighted in orange) with 13B and potential clashing BG505 residues (highlighted in pink) modeled with 13B.

Based on our structural analyses, we generated a series of mutant BG505 SOSIPs, including switching all BG505 clash residues, as well as non-clashing residues within the antibody binding footprint to B41 residues. We aimed to transfer B41-specific nAb binding properties to the BG505 trimer by generating the following changes: K232T, P240T, K347A, Q348K and K351E. The K232T, Q348K and K351E changes were included based on the above consideration that these should remove clashes with nAb 13B, while P240T and K347A were included to restore contact residues and recover binding. We note that some of the clashes may be indirect consequences of the presence of different neighboring amino acids that cause a rearrangement of peptide backbone nearby, particularly the loops containing residues 231-232 and 268-269. Therefore, we expanded the mutations to 4 different regions, including 229-232 (mut1), 350-356 (mut2), 347-348(mut3), and 240-241(N241 knock-in), while retaining the rest of the residues (FIG 6A).

**FIG 6.**
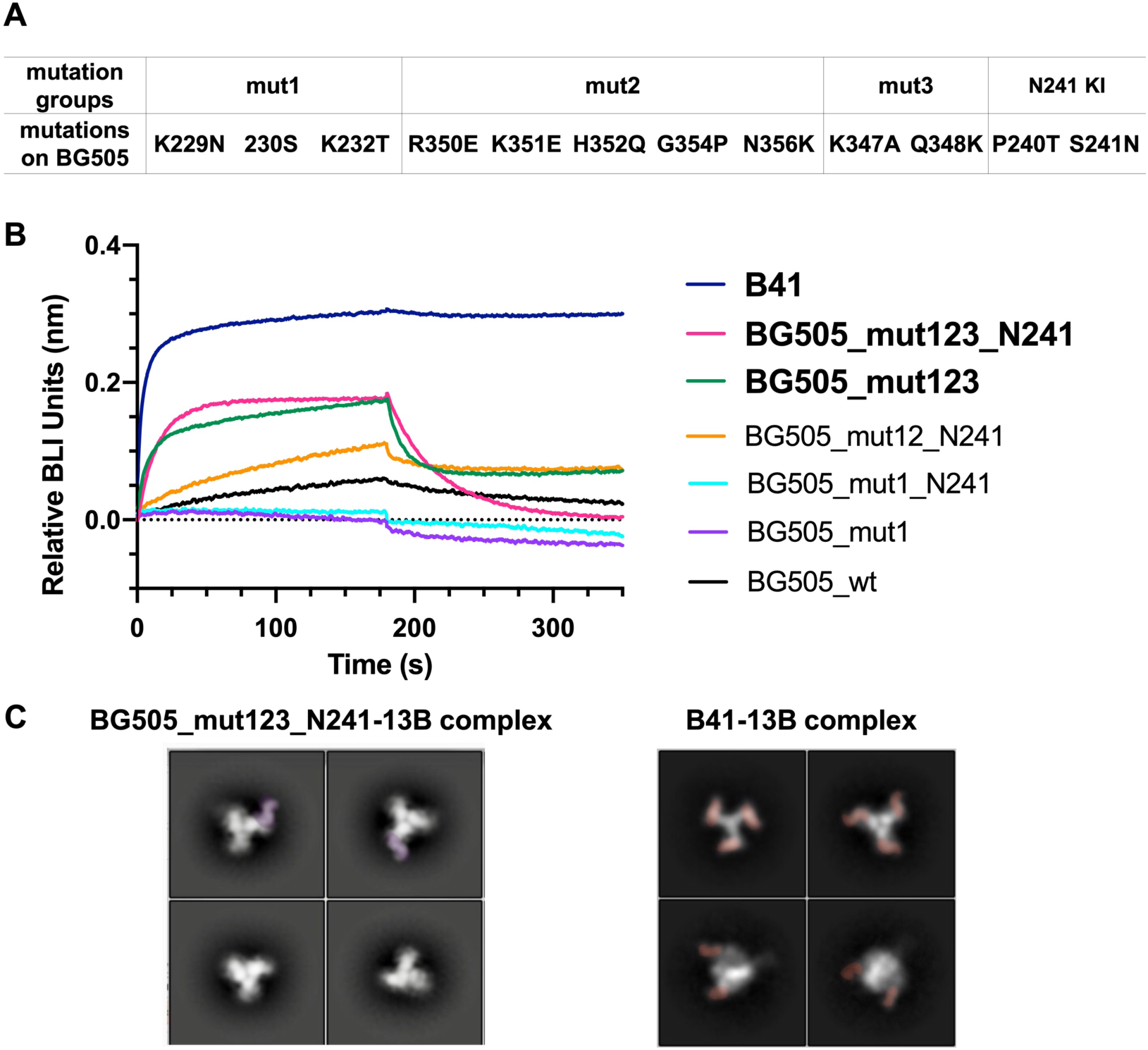
Restoration of binding of B41-specific antibody to BG505 mutants. (A) Mutations in the 4 different regions. (B) BLI binding analysis of B41 (dark blue) and a series of BG505 mutants (color codes shown on the right) against the 13B antibody. (C) Comparison of 2D classes between BG505_mut123_N241-13B complex (stoichiometry of zero to one antibodies per BG505_mut123_N241 trimer) and B41-13B complex (stoichiometry of three antibodies per B41 trimer).

BLI and NS-EM were used to screen the effect that different mutations in BG505 SOSIP on 13B binding. By combining 229-232 (mut1), 350-356 (mut2), 347-348(mut3), and 240-241(N241 knock-in) mutations (FIG 6A), we partially conferred 13B binding capabilities on BG505 SOSIP. B41 SOSIP showed strong binding and no off-rate against 13B (dark blue), whereas BG505_mut123_N241 and BG505_mut123 both exhibited binding with high off-rates against 13B, while all the other mutants exhibited similarly poor binding as 13B to BG505_wt (FIG 6B). To confirm BG505 mut123_N241 binding to 13B and obtain more structural insights into the complex, we incubated 10-fold excess Fab (molar ratio to trimer) with BG505_mut123_N241 overnight and conducted NS-EM studies. The 2D classes showed a stoichiometry of zero to one antibody per trimer (FIG 6C). Among the particles collected, ∼57% were trimers that had no Fab bound, and ∼43% of the particles had one Fab bound. No classes with more than one 13B antibody bound were found in 2D classifications even with 10-fold excess Fab. This result suggests weaker binding of BG505_mut123_N241 and 13B compared to B41 SOSIP with 13B.

To further assess the role of N241 in epitope recognition, we compared the BG505_mut123 with and without the N241 knock-in by both BLI and NS-EM analysis. The BLI results showed that, in the presence (FIG 6B, pink) and absence (FIG 6B, green) of the N241 glycan, BG505 bound 13B similarly. Although the sample lacking N241 results in slightly faster on and off rates, the overall trace is very similar, again confirming that N241 is not a crucial factor for accessing and binding to the epitope.

The NS-EM and BLI binding data both show that we enabled binding of BG505 mut123 and BG505 mut123_N241 to 13B. The relatively rapid on-rates to these mutants indicate that residues hindering the BG505 and 13B interaction have been removed. However, the high off-rate suggests that the complex is still not as stable as 13B in complex with B41 SOSIP, likely due to fewer productive interactions.

We generated some level of cross reactivity with glycan hole targeting antibody 13B by introducing B41 mutations to the glycan hole epitope region of BG505 (FIG 6). Binding of B41 and BG505 mutants with four other B41 specific glycan hole nAbs was tested by BLI (FIG 7) to ascertain whether these mutations would confer cross-reactivity to other B41 specific glycan hole antibodies. All four antibodies bound B41 as expected with binding quickly reaching saturation levels with no detectable off-rates. 16D did however exhibit a slower binding on-rate compared to others which also shows magnitude of lower neutralization against B41 pseudovirus (FIG 1D). In contrast, none of the four antibodies bound to any of the four trimer variants tested: BG505_wt, BG505_293S_deglyco, BG505_mut123, or BG505 mut123_N241. In addition to being derived from distinct antibody lineages, the sequence alignment of CDRH3 region shows that the sequence and length varies between rabbit nAbs, which likely results in different molecular interactions at the epitope-paratope interface as those observed in 13B. This result further emphasizes the strain-specific nature of the mAb responses and the different ways that antibodies can recognize the glycan hole epitope.

**FIG 7.**
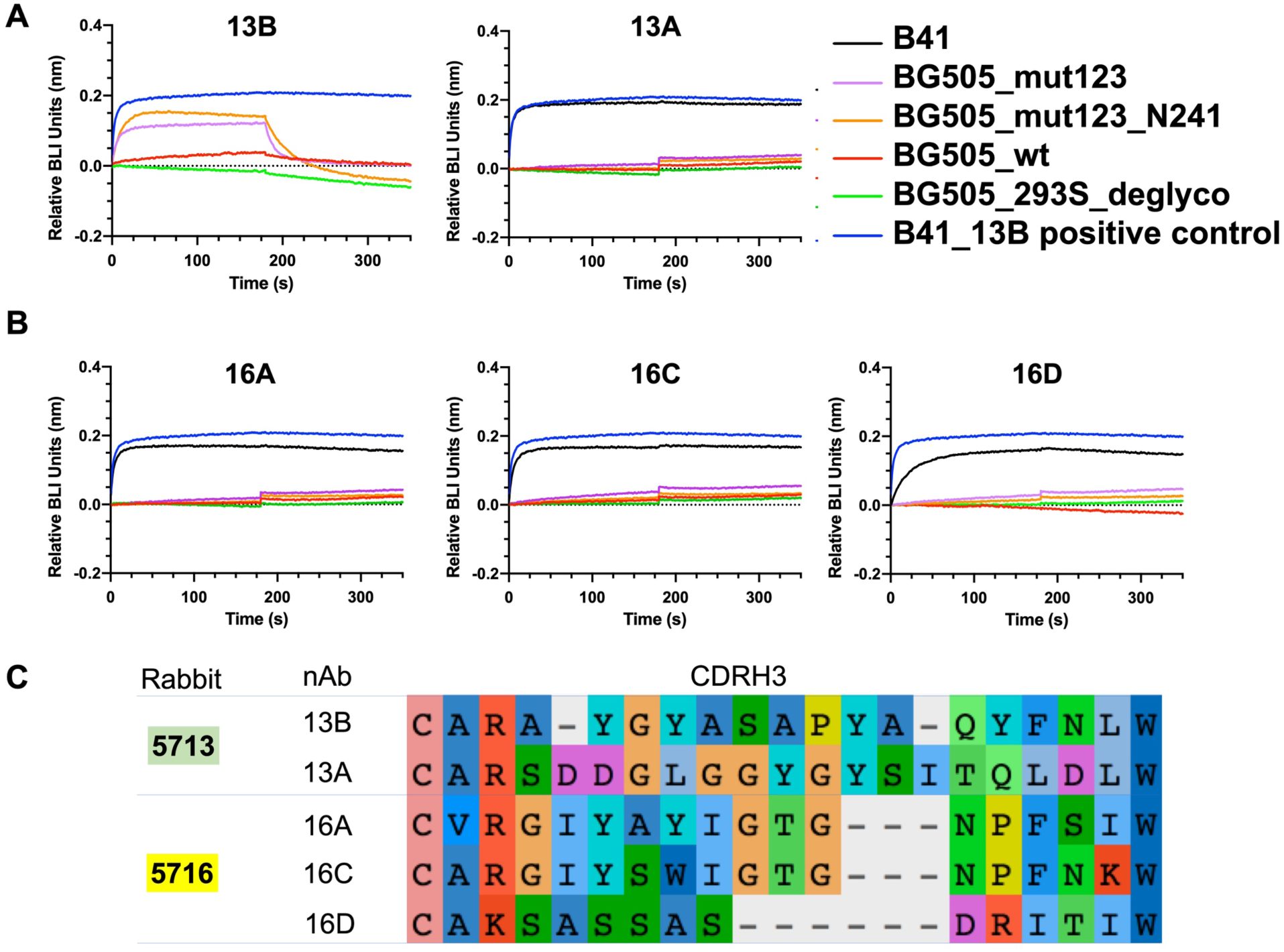
BLI analysis of B41 (black) and a series of BG505 mutants against rabbit antibodies, including(A) 13B, 13A and (B) 16A, 16C, 16D. BG505_wt (red), BG505_deglyco (green), BG505_mut123 (purple), BG505_mut123_N241 (orange), and B41_13B (blue) as positive control. (C) Sequence alignment of CDRH3 of representative rabbit nAbs. Residues are colored with default settings in AliView (22).

## DISCUSSION

Our structural studies show that immunization with the clade B trimer B41 SOSIP resulted in two epitope regions targeted by all isolated antibodies: autologous glycan hole targeting antibodies as well as non-neutralizing base-binding antibodies. Serum analyses in previous studies have shown that the glycan hole epitope region commonly elicited autologous nAb responses in B41 and BG505 SOSIP immunizations (13, 14, 18). Furthermore, polyclonal epitope analysis in rabbits (18) and non-human-primates (Ward lab, under review) of BG505 SOSIP trimer immunizations demonstrated that glycan hole antibodies are frequently elicited, making it an important epitope to understand, so that it could potentially be exploited for cross-reactive immunization strategies.

Here we endeavored to elucidate the basis for the lack of cross-reactivity of the glycan hole nAbs that were elicited by B41 and BG505 SOSIP immunogens. A comparison of the low-resolution NS-EM reconstructions of BG505 trimers in complex with BG505 nAb 11A, and B41 trimers in complex with B41 nAbs 13B and 49A revealed highly overlapping epitopes and angles of approach. While introduction of glycans at positions 230 and 289 in BG505 completely abolished neutralizing activity for 10A, 11A, and 11B (13), neutralization for the B41 nAbs was abolished for N289 but only partially reduced by ∼20% for N230, respectively, when introduced into B41. Interestingly, neutralization for N230 is increased, and even restored to some degree for N289, when the virus was produced in the presence of kifunensine, which results in more homogeneous, oligomannose glycans. This suggests that complex glycans at N289 are a particularly strong barrier to the glycan hole epitope. Removal of glycan N241 in B41 reduced neutralizing activity of the B41 nAbs, suggesting a potential direct role in binding. Our structural studies showed that 13B is in close proximity of the N241 glycan, we did not observe any specific interactions, despite a ∼70-fold reduction in neutralization when this glycan was removed. These data suggest that the absence of N241 glycan has indirect effect, for example by altering glycan processing of adjacent glycans. Altogether, these observations suggest that the B41 nAbs are impacted by more heterogeneous complex glycans, particularly when N289 is introduced. Similar to BG505 the N289 glycan knocks out neutralization of B41 glycan hole nAbs. B41 is however less impacted by the N230 glycan. Because the glycans only reduce binding and there are no specific contacts with any glycans the underlying amino acids that comprise the epitope must be responsible for the strain specificity.

The NS-EM and ELISA competition assay demonstrated that all antibodies bind to the 230/289 glycan hole epitope in a similar fashion, suggesting subtle differences at the amino-acid level. To further investigate these details, we tested the binding of B41-specific nAbs to BG505 with B41 mutants that were based on the amino-acid contacts that we observed in our cryoEM structure of B41 SOSIP bound to 13B. Among all of the different B41 nAbs tested, only 13B recovered partial binding to the BG505-mut123. This finding is not that surprising given that the sequences of CDRH3 in the B41-specific nAbs were relatively diverse. These data, and the lack of any neutralization in the viruses lacking the N289 glycan in the 117-virus panel, demonstrate the very narrow strain specificity of these mAbs (FIG S4).

We attempted to reveal the structural basis for lack of cross-reactivity nAbs that target a similar epitope region in the BG505 and B41 immunogens. Using the cryoEM structure of the B41 SOSIP trimer in complex with nAb 13B, we determined that B41 and BG505-specific nAbs target amino acids that differ between the strains. While we could recover partial binding by substituting residues, broadening antibody responses to the N289 glycan hole site is likely to remain a challenging prospect. While we could envision broadening B41-specific responses using one of our intermediate mutated trimers (e.g. BG505_mut123_N241) as a boost immunogen to bridge toward BG505 cross-reactivity, this boost would likely be specific to a single antibody lineage, namely 13B, which was only present in one rabbit. Thus, even with the increased knowledge gained from all of the analyses here, HIV immunogen design for broader antibody responses at glycan hole sites remains challenging.

## METHODS

### Immunizations

Immunization details are summarized in Fig. 1A. Animals 5713 and 5716 received 30 µg B41 SOSIP trimer alone. Animal 5746 received a BG505 SOSIP and B41 SOSIP cocktail (1 to 1 ratio) with total dose of 10 µg. Animal 5749 received a BG505 SOSIP and B41 SOSIP cocktail (1 to 1 ratio) with a dose of 30µg each time, respectively. The immunization of animals 5745 and 5748 were described in a previous study (14).

### Neutralization assays and pseudovirus production

Single-round infectious HIV-1 Env pseudoviruses were produced as described previously (Seaman et al., 2010). Briefly, plasmids encoding Env were cotransfected with an Env-deficient backbone plasmid (pSG3DENV) using Fugene 6 (Promega). Virus-containing supernatants were harvested 48 hr post-transfection, stored at −80**°**C, and then titrated on TZM-bl target cells to determine the dilution appropriate for the neutralization assays. Pseudovirus neutralization assays using TZM-bl target cells were carried out as previously described (23). Prior to evaluation, mAbs were purified as described below and passed through a 0.22 μM filter. Plasma samples were heat-inactivated at 50°C for 30 minutes and then passed through a 0.22 μM filter. mAbs and/or plasma were then serially diluted in a 96-well plate and incubated with virus for 1h prior to addition of TZM-bl target cells. After 48 hours, the relative light units (RLU) for each well were measured and neutralization calculated as the decrease in RLU relative to virus-only control wells. ID_50_/IC_50_ values are reported as the reciprocal dilution/antibody concentration that resulted in 50% virus neutralization after fitting the curve of log concentration (plasma/mAb) versus percent neutralization in Prism. For kif-grown viruses, 25 mM kifunensine was added to 293T cells on the day of transfection.

### Antibody isolation

Cryopreserved PBMCs were thawed, resuspended in 10 ml of RPMI 10% FCS and collected by centrifugation at 600 x g for 5 min. Cells were washed with PBS and resuspended in 10 ml of PBS and collected by a second centrifugation step. Cells were resuspended in 100 μl of FWB (2% FCS PBS) with anti-rabbit IgG FITC (1:1000), 1 μl of a streptavidin-PE tetramer of biotinylated BG505 SOSIP and 1 μl of a streptavidin-APC tetramer of biotinylated B41 SOSIP. After 1 h on ice, cells were washed once with 10 ml of PBS, collected by centrifugation, and resuspended in 500 μl of FWB for sorting on a BD FACS Aria III. IgG^+^ lymphocytes that stained positive for either/both BG505 or B41 tetramers were collected at 1 cell per well into Superscript III Reverse Transcriptase lysis buffer (Invitrogen) as previously described and immediately stored at −80°C.

cDNA was generated using Superscript III Reverse Transcription (Invitrogen) as previously described (24). First round PCR products were produced using 2.5 μl of cDNA and Hotstart Taq Master mix (Qiagen) for 50 cycles using the first-round primers as reported previously (13). Subsequently, 2.5 μl of first round PCR product was used as template for the second round using the second-round primers as reported previously (13). PCR products were sequenced and then analyzed using the IMGT Vquest tool. mAb lineages were identified as those with highly similar CDRH3 loop sequences. Heavy and light chain variable regions were then amplified by PCR with primers (McCoy et al.) containing homology arms specific for the expression vector. PCR products and vector were ligated using high fidelity assembly mix (NEB) into expression plasmids adapted from the pFUSE-rIgG-Fc and pFUSE2-CLIg-rK2 vectors (Invivogen). Human and rabbit Abs were transiently expressed with the FreeStyle 293 Expression System (Invitrogen). Abs were purified using affinity chromatography (Protein A Sepharose Fast Flow, GE Healthcare).

Two non-nAbs (45A and 48A) were isolated in parallel from studies described previously (14) and used as a control. The negative control mAb, named hybrid, was made from the heavy chain of R56 and light chain of R20 (PDB: 4JO3) used in a previous study (13).

### Competition ELISAs

96-well plates were coated overnight at 4°C with mouse anti-Avi-tag antibody (Genscript) at 2 μg/ml in PBS. Plates were washed 4 times with PBS, 0.05% (v/v) Tween, and blocked with 3% (w/v) BSA PBS for 1 h. Concurrently, 5-fold serial dilutions of rabbit or human mAbs were pre-incubated with 1 μg/ml of purified Avi-tagged SOSIP protein for 1 h. The mAb-SOSIP mixture was then transferred to the ELISA plate and incubated for 1 h. Plates were washed four times and incubated with 0.5 μg/ml of biotinylated mAb for 1 h, washed again, and binding detected with streptavidin-alkaline phosphatase (Jackson Immunoresearch) at 1:1000 for 1 h. mAbs were biotinylated using the NHS-micro biotinylation kit (Pierce).

### Mutations, protein expression and purification

To produce mutant viruses, the parental Env-encoding plasmid was altered by site-directed mutagenesis using the QuikChange site-directed mutagenesis kit (Agilent) according to the manufacturer’s instructions. Sanger sequencing was performed to verify that each plasmid encoded the desired mutation. Mutant pseudoviruses were then produced by co-transfection with pSG3DENV as described above. Mutations in BG505 SOSIP were generated by Agilent QuikChange Lightning Multi Site-Directed Mutagenesis Kit and confirmed by Genewiz sequencing.

All untagged B41 SOSIP and BG505 SOSIP were expressed using HEK 293F cells for 6 days and then purified on a 2G12 IgG cross-linked Sepharose column. The proteins were eluted by 3 M MgCl_2_, pH 7.2 buffer, and then further purified over a HiLoad 16/600 Superdex 200 pg column in 20 mM Tris pH 7.4, 150 mM NaCl (1x TBS) buffer.

All C-term His_6_-tagged B41 SOSIP and BG505 SOSIP mutants were expressed using HEK 293F cells for 6 days and then purified on a 2G12 IgG cross-linked Sepharose column. The proteins were eluted by 3 M MgCl_2_, pH 7.2 + 250 mM L-Arginine buffer and then further purified over a HiLoad 16/600 Superdex 200 pg column in 20 mM Tris pH 7.4, 150 mM NaCl + 250 mM L-Arginine buffer. 250 mM L-arginine waswas added to prevent aggregation for Env trimers with added C-term His_6_-tags.

BG505 with high mannose glycans was expressed using HEK 293S cells for 6 days and then purified on a 2G12 IgG cross-linked Sepharose column. The trimers were eluted by 3M MgCl_2_, pH 7.2 + 250 mM L-Arginine buffer and then further purified over a HiLoad 16/600 Superdex 200 pg column in 20 mM Tris pH 7.4, 150 mM NaCl + 250 mM L-Arginine buffer. The purified BG505 was then cleaved by EndoH enzyme overnight to remove glycans and then purified over a HiLoad 16/600 Superdex 200 pg column in 20 mM Tris pH 7.4, 150 mM NaCl + 250 mM L-Arginine buffer.

Fabs from rabbits were expressed in 293F cells for 6 days and then affinity purified using a CaptureSelect™ CH1-XL Pre-packed 1 ml Column (ThermoFisher).

### Biolayer interferometry

His-tagged B41 SOSIP and His-tagged BG505 SOSIP variants at 0.05 mg/ml were loaded onto Ni-NTA biosensors and dipped into 1 µM (300 µl) of rabbit Fab using an Octet Red96 instrument (ForteBio). After loading for 180 s, association was measured for 180 s followed by dissociation for 600 s in 1 X kinetics buffer (phosphate-buffered saline pH 7.2, 0.01% [w/v] BSA, 0.002% [v/v] Tween-20). A baseline containing no trimer sample, but the same concentration of Fab in 1 X kinetics buffer, was subtracted from each data set and curves were aligned on the Y-axis using the baseline step. Baseline subtraction minimized influence of non-specific binding of Fab to the sensor tip.

### Negative-stain EM sample preparation, data collection and processing

All trimer-Fab complexes were generated by incubating 10X molar Fab with B41 SOSIP or BG505 SOSIP mutants overnight at room temperature. Grid preparation, image processing, and raw data analysis followed a similar protocol described previously (25). Briefly, samples were diluted with 1x TBS to 0.01 mg/ml right before putting on grids. Three µl of sample was then applied to a 400 mesh carbon-coated Cu grid, then stained with 2% (w/v) uranyl formate for 45-60 s. Grids were blotted using blotting paper untilcompletely dry. All grids were imaged on a 120 keV FEI Tecnai Spirit electron microscope using a nominal magnification of 52000x, resulting in 2.05 Å/px. Micrographs were collected with a TVIPS TemCam-F416 (4k x 4k) camera using the Leginon interface (26) with a defocus of 1.5 μm.

Particles were selected using Appion DoGPicker (27) and extracted with Relion v2.1.(28) Extracted particles were imported to cryosparc v2.8.0 (29). Particles were then classified in 2D into 50 classes. Classes not containing features of trimers were removed, and the remaining particles were used for 3D refinement. The NS-EM 3D reconstructions have been deposited to the Electron Microscopy Data Bank: B41-45A (EMD-20882), B41-48A (EMD-20737), and B41-49A (EMD-20738).

### Cryo-EM sample preparation, data collection and processing

B41 SOSIP trimers were complexed with 6X molar excess of 13B Fab overnight at room temperature and then purified by HiLoad 16/600 Superdex 200 pg column column. The eluted sample were concentrated to 5 mg/ml and applied to previously plasma-cleaned Protochips C-flat 2/2 400 mesh Cu grids and blotted once for 5 s with blot force 0 after a wait time of 10 s. Blotted grids were plunge frozen into nitrogen-cooled liquid ethane using a Vitrobot Mark IV (ThermoFisher).

Data were collected on a Talos Arctica operating at 200 kV coupled with a K2 Summit direct electron detector at a nominal magnification of 36000x resulting in 1.15 Å pixel size. Dose was calculated to be 5.67 e^-^/pix/s. 47 frames were collected per movie with 250 ms exposure time each, resulting in a total dose of ∼50.4 electrons Å^−2^. Micrographs were collected with the automated Leginon interface (26) using a defocus range from 0.8 to 2 μm.

Movies were aligned and dose-weighted using MotionCor2 (30) (FIG S1A). Data were then processed with Cryosparc v2 (29) (FIG S1B). A total of 1,721 micrographs were used. Particles were then classified in 2D by 50 classes, and classes not containing features of trimers were removed, resulting in particle images that were retained for further processing.

The final refinement included 145,000 particles and C3 symmetry was imposed. The resolution of the final map was calculated to ∼ 3.9 Å at a Fourier shell correlation (FSC) cut-off at 0.143. The EM reconstructions have been deposited to the Electron Microscopy Data Bank (EMD-20642).

### Modeling and refinement of cryo-EM structures

Initial homology models of B41 (gp120, and gp41) were generated using the crystal structure of B41 (PDB: 6MCO). An initial model of the Fv region of 13B was generated using the Rosetta antibody protocol available on the ROSIE server (20). Individual chains were fit into the 3.9 Å cryo-EM map using UCSF Chimera (21). Glycans were built with Coot (31). Sugar molecules with disordered or no density were removed. The structure was then refined using Rosetta real space refinement (32), requesting an output of 319 refined structural models. The top scoring structure was chosen after evaluation with MolProbity (33), EMRinger (34), and manual inspection. The model was iteratively refined using Rosetta real space refine and improved by manual inspection using Coot. Cryo-EM data collection and refinement statistics are summarized in Table S1. Structural figures were made in UCSF Chimera. Regions with relatively poor density in the model were removed. The model has been deposited to the Protein Data Bank (PDB: 6U59).

## Supplementary Information

**FIG S1.**
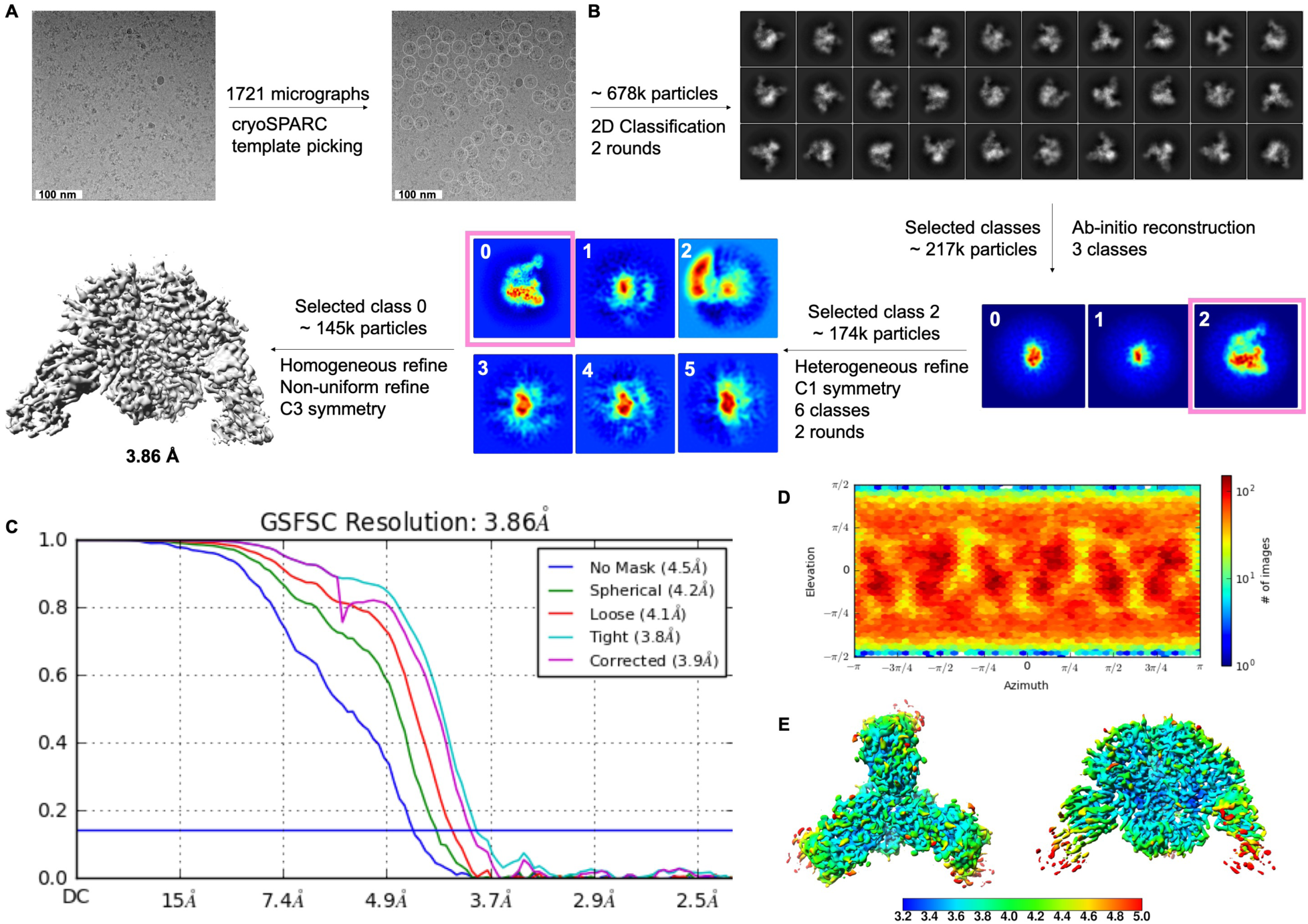
CryoEM data processing for the B41-13B complex. (A) Representative aligned and dose-weighted cryo-EM micrograph of B41-13B complex in vitreous ice, circles showing particles picked up by cryosparc v2. template picking. (B) Cryo-EM data processing scheme to obtain final reconstruction. (C) FSC plots of unmasked (blue) and masked (pink) reconstructions. (D) Relative angular distribution of final reconstruction used for model building. Red bars represent views with more particles. (E) Cryo-EM map colored according to local resolution determined by local resolution function in cryoSPARC v2. Color key shows local resolution in Å.

**FIG S2.**
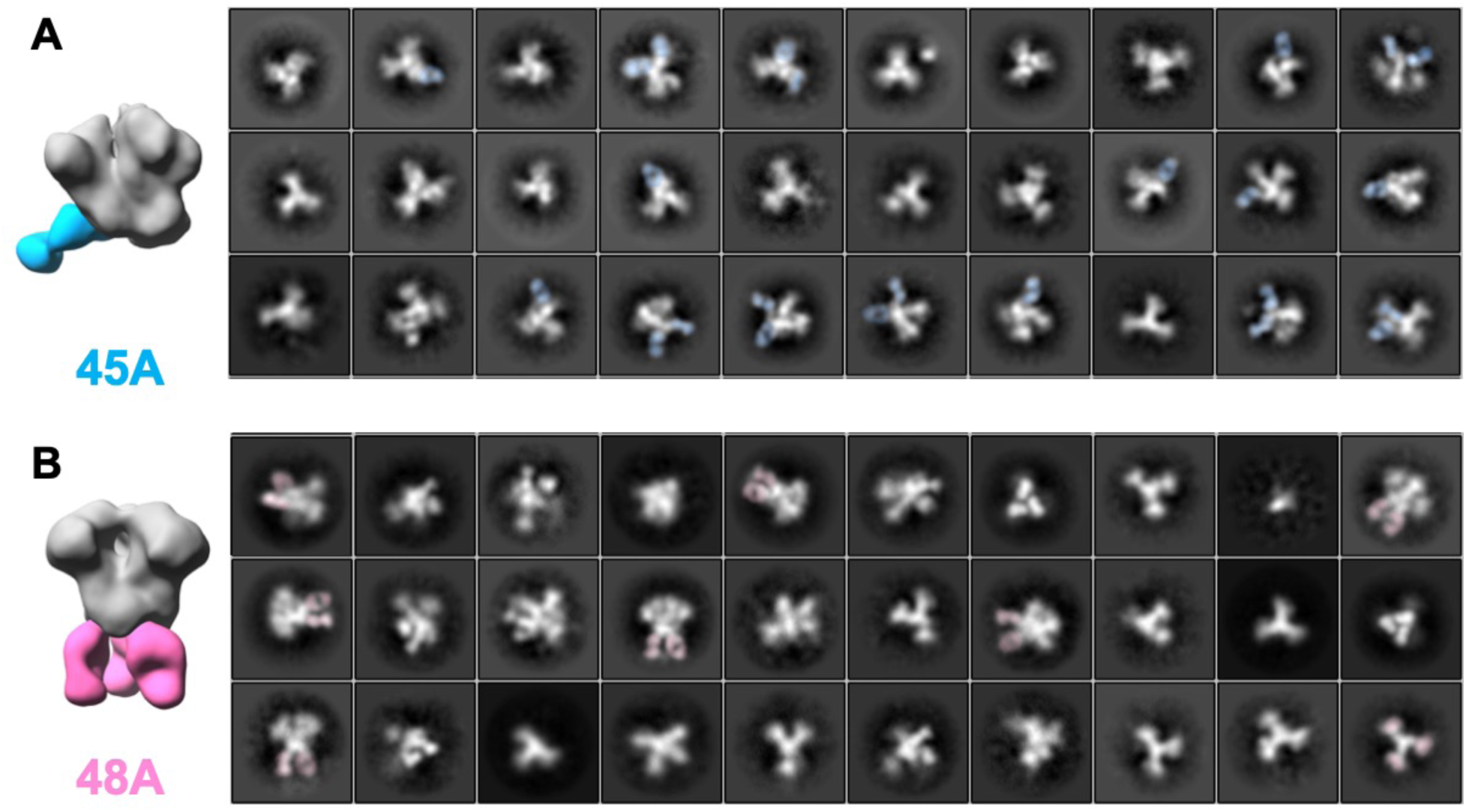
Negative stain EM 2D classification using cryoSPARC. Trimers are shown in complex with base binding antibodies 45A (blue) or 48A (pink). Classes that resemble complexes were selected for 3D reconstruction (left panel), C1 symmetry is applied for B41-45A complex and C3 is applied for B41-48A complex.

**FIG S3.**
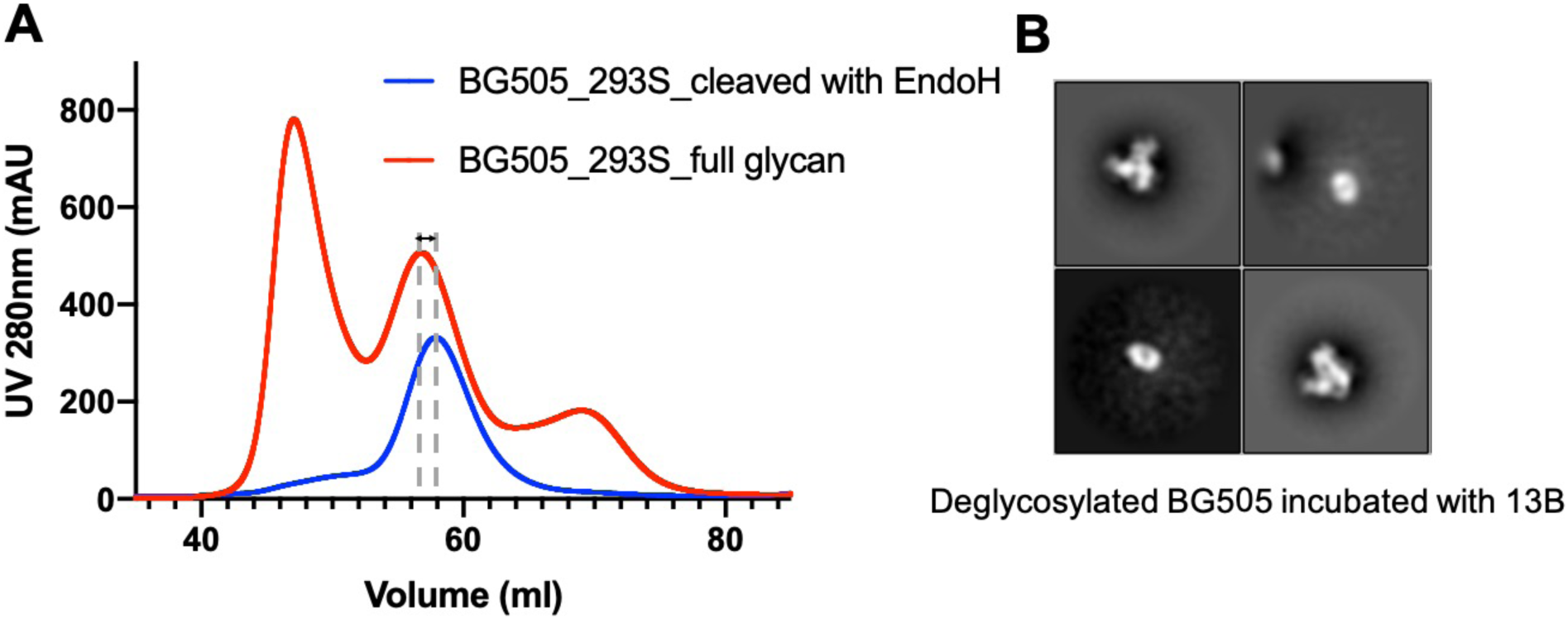
Deglycosylated BG505 characterization. (A) SEC characterization of deglycosylated BG505. HEK 293S expressed BG505 was purified over a 2G12 column and then by SEC (red line). The SEC-purified sample was then treated with EndoH overnight and then purified through the same column (blue). (B) Representative NS-EM 2D classes of 10-fold excess 13B antibody incubated with deglycosylated BG505 overnight at RT.

**FIG S4.**
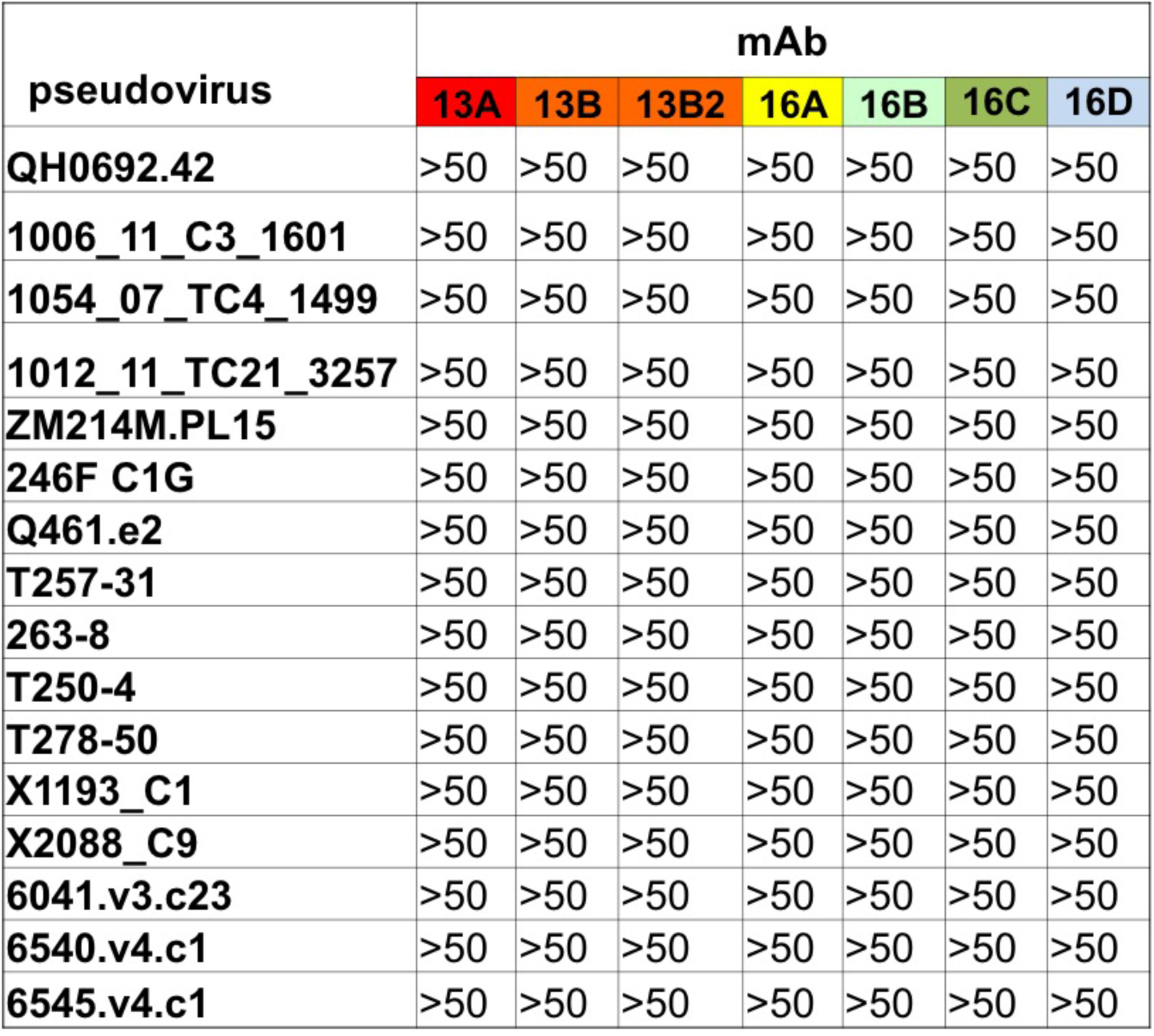
Neutralization analysis of selected viruses lacking glycan sites at 289 from the 117-virus panel against B41-specific mAbs.

**FIG S5.**
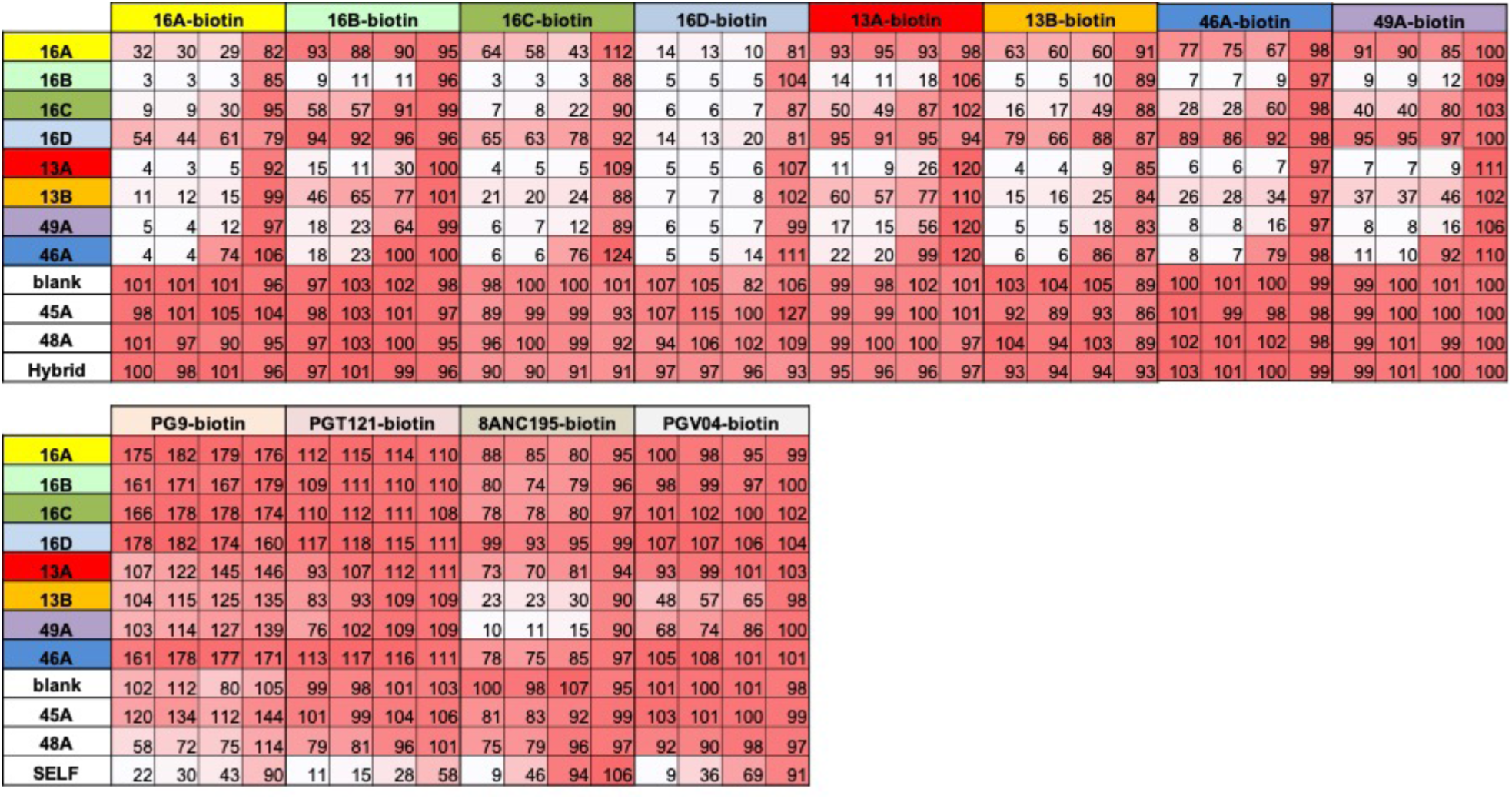
Competition ELISAs of B41-spsecific mAbs and human bnAbs PG9, PGT121, 8ANC195, and PGV04. Competition is expressed as percentage binding where 100% was the absorbance measured when B41 SOSIP protein only was captured on the anti-avi-tag ELISA plate. The non-HIV specific mAb hybrid was used as a negative control for non-specific inhibition of biotinylated rabbit mAb binding. The unbiotinylated version of each human bnAb (PG9, PGV04, PGT121, 8ANC195) was used as a postive control (referred to as SELF) only in the assay where the binding of each of these bnAbs in biotinylated form was being assessed.

**Table S1.**
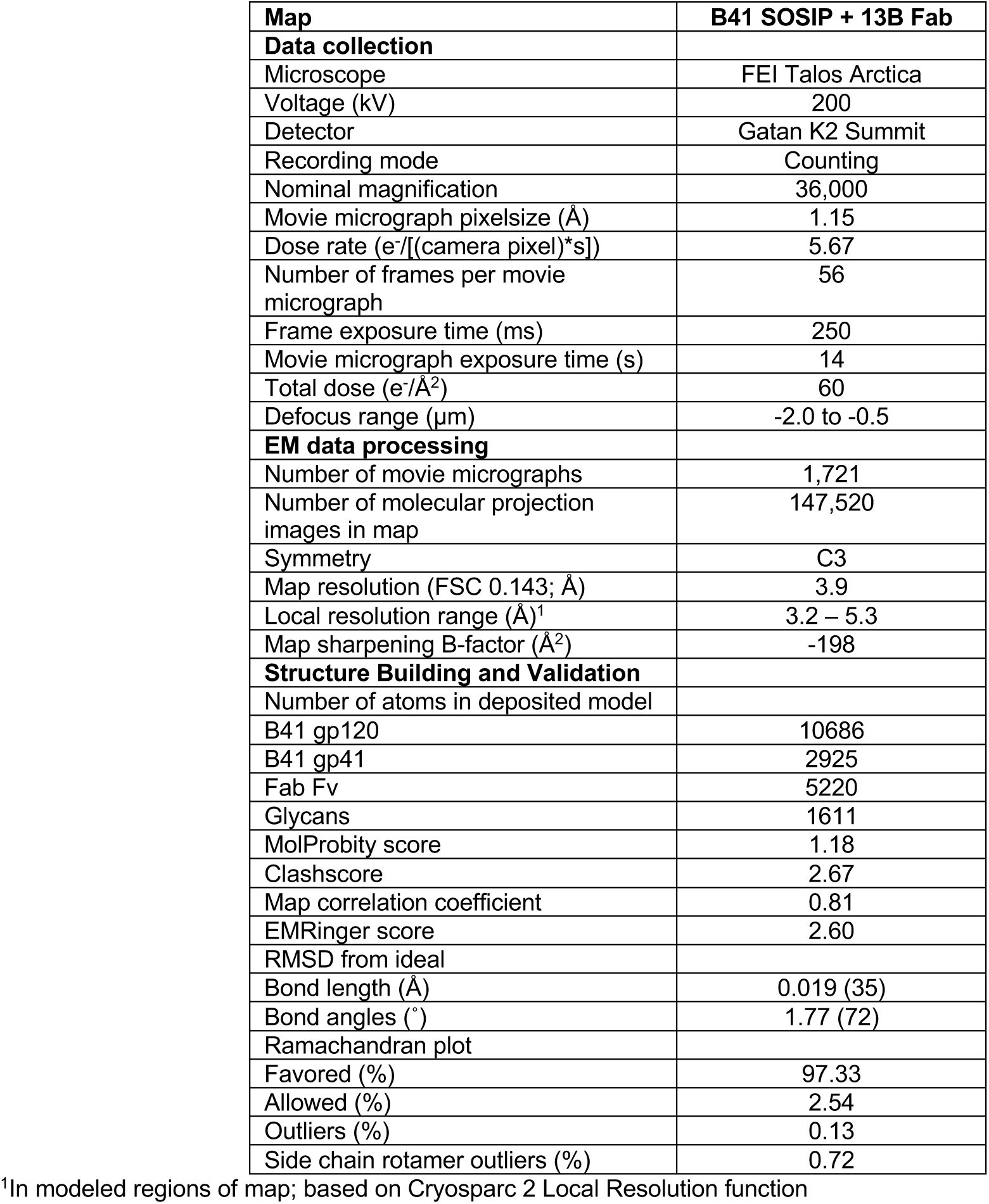
Cryo-EM data collection and refinement statistics.

| Map | B41 SOSIP + 13B Fab |
| --- | --- |
| **Data collection** |  |
| Microscope | FEI Talos Arctica |
| Voltage (kV) | 200 |
| Detector | Gatan K2 Summit |
| Recording mode | Counting |
| Nominal magnification | 36,000 |
| Movie micrograph pixelsize (Å) | 1.15 |
| Dose rate (e<sup>-</sup>/[(camera pixel)*s]) | 5.67 |
| Number of frames per movie micrograph | 56 |
| Frame exposure time (ms) | 250 |
| Movie micrograph exposure time (s) | 14 |
| Total dose (e<sup>-</sup>/Å<sup>2</sup>) | 60 |
| Defocus range (µm) | -2.0 to -0.5 |
| **EM data processing** |  |
| Number of movie micrographs | 1,721 |
| Number of molecular projection images in map | 147,520 |
| Symmetry | C3 |
| Map resolution (FSC 0.143; Å) | 3.9 |
| Local resolution range (Å)<sup>1</sup> | 3.2 – 5.3 |
| Map sharpening B-factor (Å<sup>2</sup>) | -198 |
| **Structure Building and Validation** |  |
| Number of atoms in deposited model |  |
| B41 gp120 | 10686 |
| B41 gp41 | 2925 |
| Fab Fv | 5220 |
| Glycans | 1611 |
| MolProbity score | 1.18 |
| Clashscore | 2.67 |
| Map correlation coefficient | 0.81 |
| EMRinger score | 2.60 |
| RMSD from ideal |  |
| Bond length (Å) | 0.019 (35) |
| Bond angles (°) | 1.77 (72) |
| Ramachandran plot |  |
| Favored (%) | 97.33 |
| Allowed (%) | 2.54 |
| Outliers (%) | 0.13 |
| Side chain rotamer outliers (%) | 0.72 |

<sup>1</sup>In modeled regions of map; based on Cryosparc 2 Local Resolution function

## ACKNOWLEDGEMENTS

The research was supported by NIH grant UM1AI100663 (I.A.W., A.B.W. and D.R.B.), UM1AI144462 (I.A.W., A.B.W. and D.R.B.) and P01 AI110657 (I.A.W., A.B.W., R.W.S.), the Bill and Melinda Gates Foundation grants OPP1115782 (A.B.W.) and OPP1132237 (R.W.S.), amfAR grant 109514-61-RKVA (M.J.G.). RWS is a recipient of a Vici grant from the Netherlands Organization for Scientific Research (N.W.O.). C.A.C. was supported by NIH F31 Ruth L. Kirschstein Predoctoral Award Al131873 and by the Achievement Rewards College Scientists Foundation.

We are grateful to Bill Anderson for expert microscopy assistance, to Lauren Holden, Aleks Antanasijevic, and Julianna Han for manuscript proofreading and editing, to Leigh Sewall and Jeffrey Copps for expert biochemical and technical assistance.

